# *Inoviridae* prophage and bacterial host dynamics during diversification, succession and Atlantic invasion of Pacific-native *Vibrio parahaemolyticus*

**DOI:** 10.1101/2023.03.23.534014

**Authors:** Randi L. Foxall, Jillian Means, Ashley L. Marcinkiewicz, Christopher Schillaci, Kristin DeRosia-Banick, Feng Xu, Jeffrey A. Hall, Stephen H. Jones, Vaughn S. Cooper, Cheryl A. Whistler

## Abstract

The epidemiology of *Vibrio parahaemolyticus,* the leading cause of seafood-borne bacterial gastroenteritis of humans world-wide, dramatically changed in the United States following the establishment of a Pacific native lineage called sequence type (ST) 36 in the Atlantic. In this study we used phylogeography based on traceback to environmental source locations and comparative genomics to identify features that promoted evolution, dispersal, and competitive dominance of ST36. The major genomic differentiation and competitive success of ST36 was associated with a striking succession of filamentous prophage in the family *Inoviridae* (inoviruses), including loss of an inovirus prophage that had been maintained for decades in the endemic north Pacific population. Subsequently, at least five distinct progenitors arising from this diversification translocated from the Pacific into the Atlantic and established four geographically defined clonal subpopulations with remarkably low migration or mixing. Founders of two prevailing Atlantic subpopulations each acquired new stable and diagnostic inoviruses while other subpopulations that apparently declined did not. Broader surveys indicate inoviruses are common and active among the global population of *V. parahaemolyticus* and though inovirus replacements, such as in ST36, appear to be infrequent, they are notable in pathogenic lineages that dispersed.

**Importance:** An understanding of the processes that contribute to emergence of pathogens from environmental reservoirs is critical as changing climate precipitates pathogen evolution and population expansion. Phylogeographic analysis of *Vibrio parahaemolyticus* hosts combined with analysis of their *Inoviridae* phage resolved ambiguities of diversification dynamics which preceded successful Atlantic invasion by the epidemiologically predominant ST36 lineage. It has been established experimentally that filamentous phage can limit host recombination, but here we show that phage loss is linked to rapid bacterial host diversification during epidemic spread in natural ecosystems alluding to a potential role for ubiquitous inoviruses in the adaptability of pathogens. This work paves the way for functional analyses to define the contribution of inoviruses in the evolutionary dynamics of environmentally transmitted pathogens.

## Introduction

Cases of *V. parahaemolyticus*, the leading cause of bacterial seafood-borne gastroenteritis of humans world-wide recently increased along the United States (US) Atlantic coast (1–3). Rising seasonal illnesses were tied to warming ocean temperatures, a growing aquaculture industry, and the emergence of endemic pathogens (3–6). But incursion of a Pacific Northwest (PNW) lineage called sequence type (ST) 36 into the Atlantic following an anomalously mild winter is the most important driver of this shift (3, 5, 7). ST36 began causing infections in 1979 (8) and was originally limited to the Pacific Northwest (PNW) (9). Although sporadic infections were occasionally reported outside the PNW, local sources were rarely implicated (7, 9, 10). This changed in 2012 when ST36 began causing illnesses traced to Atlantic sources and in 2013 it caused a 13-state outbreak traced to multiple northeast US locations (5, 7). Unlike previous non-endemic strains causing outbreaks from Atlantic sources, including the pandemic ST3 (11), and ST8 (12), ST36 continues to cause sporadic disease from a few northeast US locations. A better understanding of how and where ST36 established populations is needed to aid in management to limit illness. Furthermore, this expansion of ST36 provides a unique opportunity to broaden our understanding of the population and environmental context by which pandemic strains arise and spread.

Spatiotemporal analyses of the epidemiology of ST36 identified that a new population arose through recombination and replaced the original PNW population by 2000 (9). Disease patterns implied potential reciprocal transfer between both US coasts, though inference was informed by a few cases where infection occurred from shellfish consumed on the opposite coast from where illnesses were reported and further complicated by modeling of a mode of spread through human populations (9), whereas seasonal illnesses by *V. parahaemolyticus* are typically vectored by environmentally contaminated raw seafood. This highlights a continuing challenge where a lack of environmental traceback in clinical reporting that uncouples strains from their source bodies of water obscures environmental distribution (5, 9). As with other human pathogens originating from the natural environment, this leaves a critical gap of knowledge of the ecological context of pathogen evolution and expansion.

Bacteriophages, the viruses that infect bacteria, can shape, and quell environmentally vectored human pathogen populations (13–19). Lytic phages alter the dynamics of bacterial competition though selective predation, sometimes of the most numerous bacteria, thereby maintaining population diversity (20–22). Phages also impact diversity through transfer of novel DNA including toxin-encoding genes and lysogenic conversion (23–28). Chromosomally integrated prophages are double-edged swords in that they can benefit their host by excluding superinfections by related phage and by attacking their hosts’ competitors, but they can also harm their hosts by diverting resources for virion production or by host death upon lytic induction (25–27, 29, 30). Lytic phages are abundant in habitats of *V. parahaemolyticus* (31), and also under consideration for detection and biological control (32–37). Though the roles of phage in mediating bacterial competition among *Vibrio cholerae* is well appreciated (14, 38), with the exception of a few descriptive reports (39–44), the contribution of prophage to *V. parahaemolyticus* population structure is undetermined.

Prior studies have produced valuable insight into the growing problem of the ST36 lineage’s increasing clinical prevalence and potential to spread globally (9, 45), but the adaptations that prompted lineage replacement and dispersal are not evident nor are the ecological factors that spurred diversification. Here, we examined the phylogenomics of the Atlantic ST36 incursion—accompanied by interactions with filamentous *Inoviridae* phage—by capturing a broad geographic and temporal distribution of the population. By curating strains with environmental traceback, we determined the phylogeography of ST36 wherein multiple strains that first diversified in the Pacific, subsequently translocated to the Atlantic, and established spatially distinct and non-mixing subpopulations therein. This also revealed that changes in inovirus content accompanied diversification and lineage replacement in the PNW and preceded clonal expansion by distinct founders of two persisting Atlantic ST36 populations that continue to cause sporadic disease.

## Results

### Multiple distinct ST36 lineages clonally expanded to form geographically stable Atlantic subpopulations

To investigate the evolution of ST36 as it translocated from the PNW to the Atlantic, we curated a collection of genomes and mapped their environmental sources onto their constructed whole genome phylogenies. Locations include two Pacific (PNW and California [CA]), and five Atlantic (Galicia Spain, the Gulf of Maine [GOM], an Island south of Cape Cod, [SCC], Long Island Sound [LIS] and the mid-Atlantic coast [MAC] [Table S1]). The phylogenies exemplify how multiple clades emerged from the old-PNW population (Fig. 1) and gave rise to the new-PNW population (9). A single relic of the old-PNW population and 103 isolates from seven lineages arising from the modern ST36 diversification were traced to environmental locations outside the PNW (Fig. 1, clades I-VII). Notably, most translocated clades either contain a PNW-traced isolate or are most closely related to clades dominated by PNW isolates suggesting these differentiated in the PNW prior to translocation rather than after arrival in the Atlantic.

**Figure 1.**
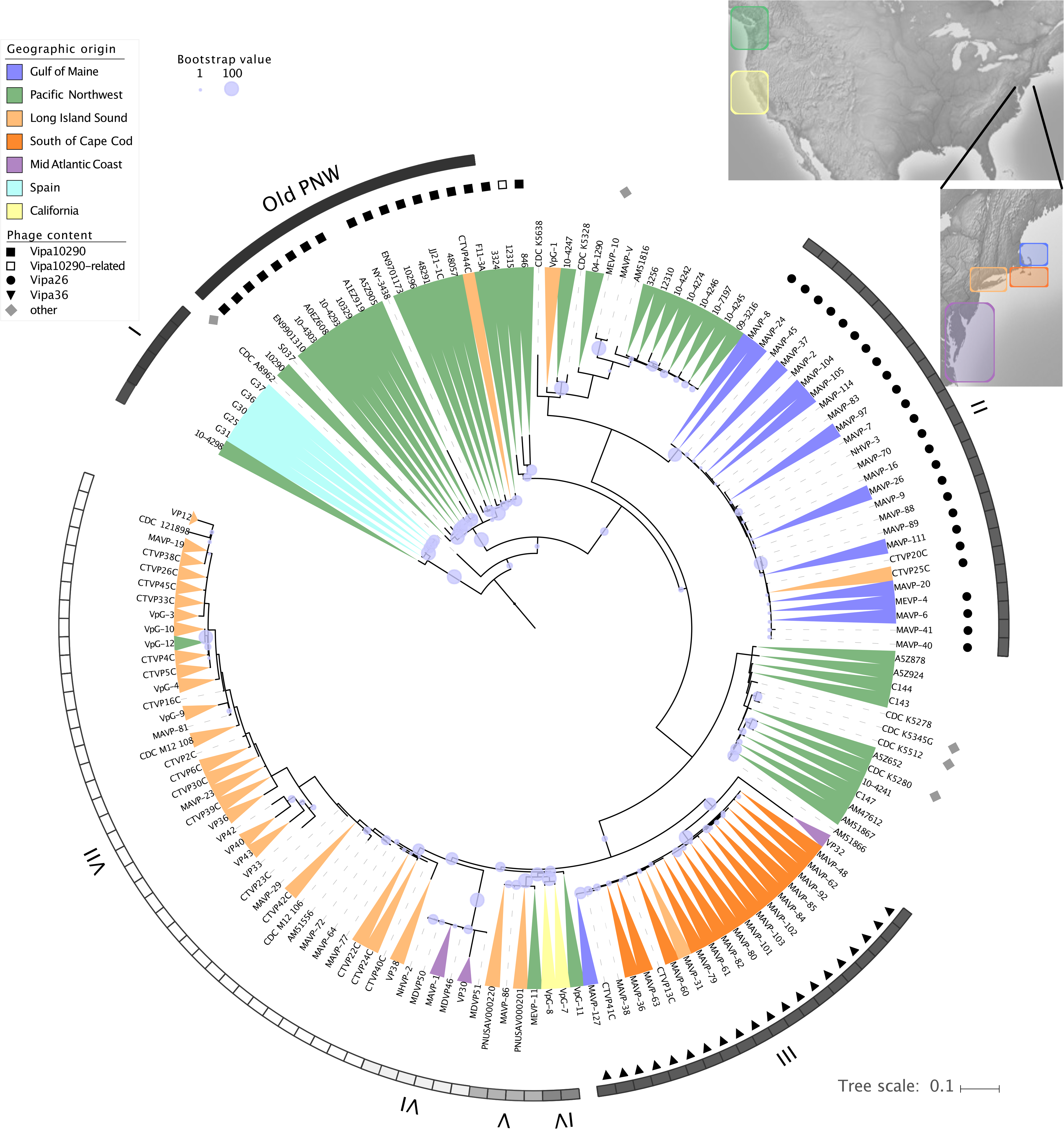
Maximum-likelihood phylogeny of ST36 *Vibrio parahaemolyticus*. ML-phylogenies were built with 1,025,281 aligned variable nucleotide sites identified in quality ST36 isolates. Isolates are colored to correspond to geographic environmental traceback, where available (Table S1) and where no color indicates unknown or ambiguous origin. Isolates acquired from four Northeast US States as part of this study are named uniquely by a combination of state (Maine [ME], New Hampshire [NH], Massachusetts [MA], and Connecticut [CT]) of reporting and sequential numbers. Symbols next to isolates indicate identity of inovirus content. The ancestral PNW ST36 population is identified by a black bar, whereas seven different lineages associated with translocation events (I – VII) are identified by greyscale bars. Bootstraps are from 250 replicates (criterion reached).

Assignment of isolates to environmental sources revealed striking geographic structure of the Atlantic subpopulations in GOM, SCC, and LIS, indicating that each was founded by a genetically unique individual (Fig. 1). All but one isolate traced to the GOM are clonal (clade II) and share ancestry with PNW isolates (9, 46–48), and the 2006 New York isolate VpG-1 (9)(Fig. 1, Table S1). Most isolates traced to other Atlantic locations and those from CA share common ancestry with isolates reported from British Columbia Canada in 2005, and later in WA in 2011-2012 (Fig. 1 and Table S1). All but one SCC-traced isolate (clade III) are clonal and only two clade members trace to other nearby locations. LIS-traced isolates are also mostly clonal (76%) (clade VII), though two distinct 2018 isolates group within a mixed-location clade (clade V) with isolates trace to the PNW in 2011, and from oysters consumed in CA in 2015. Phylogenies built with core non-recombining variation mostly agreed with whole genome phylogenies, though a change in clonal assignment of two strains (CTVP25C and MAVP-48) alluded to the possibility that horizontally acquired variation these shared in common with the GOM and SCC clades respectively may have obscured their distinct heritage (Fig. 1, S1 & S2; Tables S2-S4).

Given the geographic linkage of clonal subpopulations (Fig. 1), we examined the population genetic structure of isolates from known locations to identify patterns of coancestry and admixture (49). The conservative use of core variation and eight ancestral populations (which best explains the data, see Fig. S3A & B) identified two old-PNW populations that were replaced by a single modern-day PNW population, comprised of two geographically distinct lineages: one related to GOM isolates and the other to most SCC/LIS/MAC isolates (Fig. 1 & Fig. 2). Six PNW isolates were exceptions in that they belong to the same population and form clades with other translocated isolates, including those traced to Spain. Although no PNW isolates belong to the population that translocated to CA, two PNW isolates exhibit a shared history of admixture with the CA isolates providing clues to this lineage’s origin. The clonal clades from SCC and LIS, which are in very close proximity to each other and without geographic barriers, also belong to the same population, whereas two MAC isolates form a distinct population. Atlantic-derived isolates from locations south of the GOM exhibit little coancestry with other populations, whereas the GOM clade displays near equal mixed coancestry between the prevalent modern PNW population and the other Atlantic populations (Fig. 2). Five PNW isolates also exhibited this same pattern of mixed heritage.

**Figure 2.**
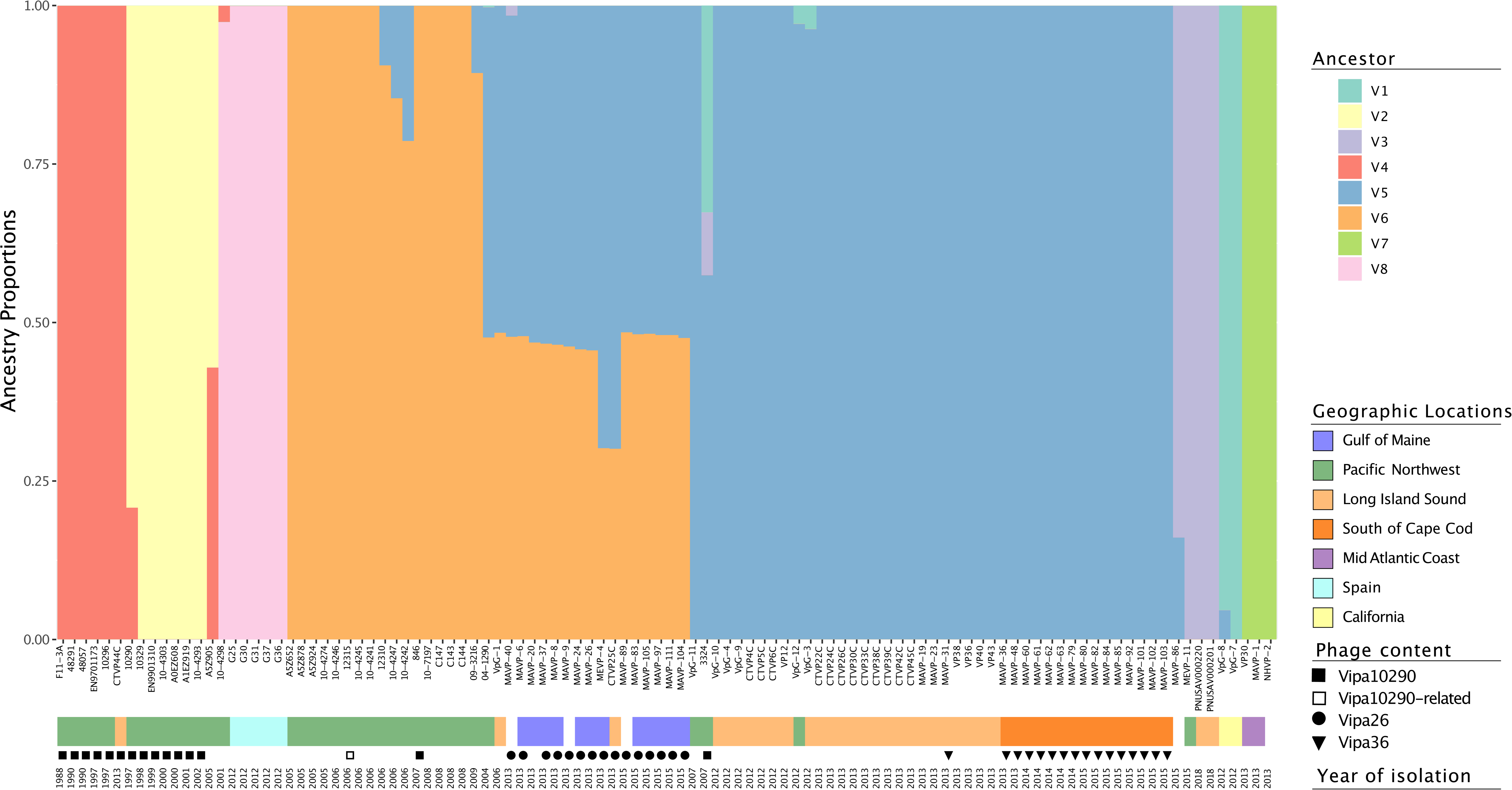
Population structure of ST36 strains. Coancestry estimates were inferred from SNP matrices using 741 SNPs identified from core non-recombining genomes of ST36 isolates with known environmental source and at least one close relative (see Table S5 for excluded genomes) by LEA (49). Colored bars represent proportion of genetic variation derived from 8 ancestral populations (see Fig S3A). Geographic locations are marked by colors below strain name, followed by phage content (symbol) and year of isolation.

Analysis of admixture that incorporated horizontally acquired content separated the two Atlantic populations on opposite sides of Cape Cod, MA (GOM and SCC) from the rest of those in the Atlantic which grouped with strains from CA (Fig. S4). This also relocated a single GOM clade isolate (MAVP-20) into the CA/MAC/LIS population. This suggested that horizontally acquired variation in the SCC and GOM lineages (that MAVP-20 lacks) may define their uniqueness. In contrast to the GOM and SCC populations that exhibit little admixture, the modern PNW ST36 isolates exhibit complex admixture as do members of the LIS population (Fig. S4). Relatively few individuals within the GOM and SCC populations display some admixture with each other. Thus, population structures inferred from horizontal variation better aligned most strains to their geographic locations even as horizontal content overshadowed some interesting ancestral patterns, specifically, the shared mixed heritage of the GOM clade with modern PNW strains (Fig. 2).

### Inovirus loss and reacquisition accompanied ST36 population replacement and expansion

One explanation for the differences in core versus whole genome population structure is that differential genetic gains or losses preceded clonal expansion. To identify such features, we compared high-quality genomes of four Atlantic clade members, (MAVP-1, MAVP-23, MAVP-36, and MAVP-26), to PNW strains of the new-PNW (12310) and old-PNW (10290) clades. This revealed MAVP-26, MAVP-36, and 10290 each contain a unique prophage integrated into the *dif* site at the replication terminus of chromosome I, similar to prophage f237 found in pandemic O3:K6 strains (39, 41) (Fig.3). In contrast, the other two Atlantic clade members and the new-PNW isolate lack a phage in this location (Fig. 3). These are classified in the family *Inoviridae* (hereafter inoviruses) and we assigned unique names, including “vB” for <u>v</u>irus infecting <u>B</u>acteria, and “Vipa” in reference to the host *<u>Vi</u>brio <u>pa</u>rahaemolyticus* and isolate name hereafter Vipa26 in MAVP-26, Vipa36 in MAVP-36, and Vipa10290 in strain 10290 (50).

**Figure 3.**
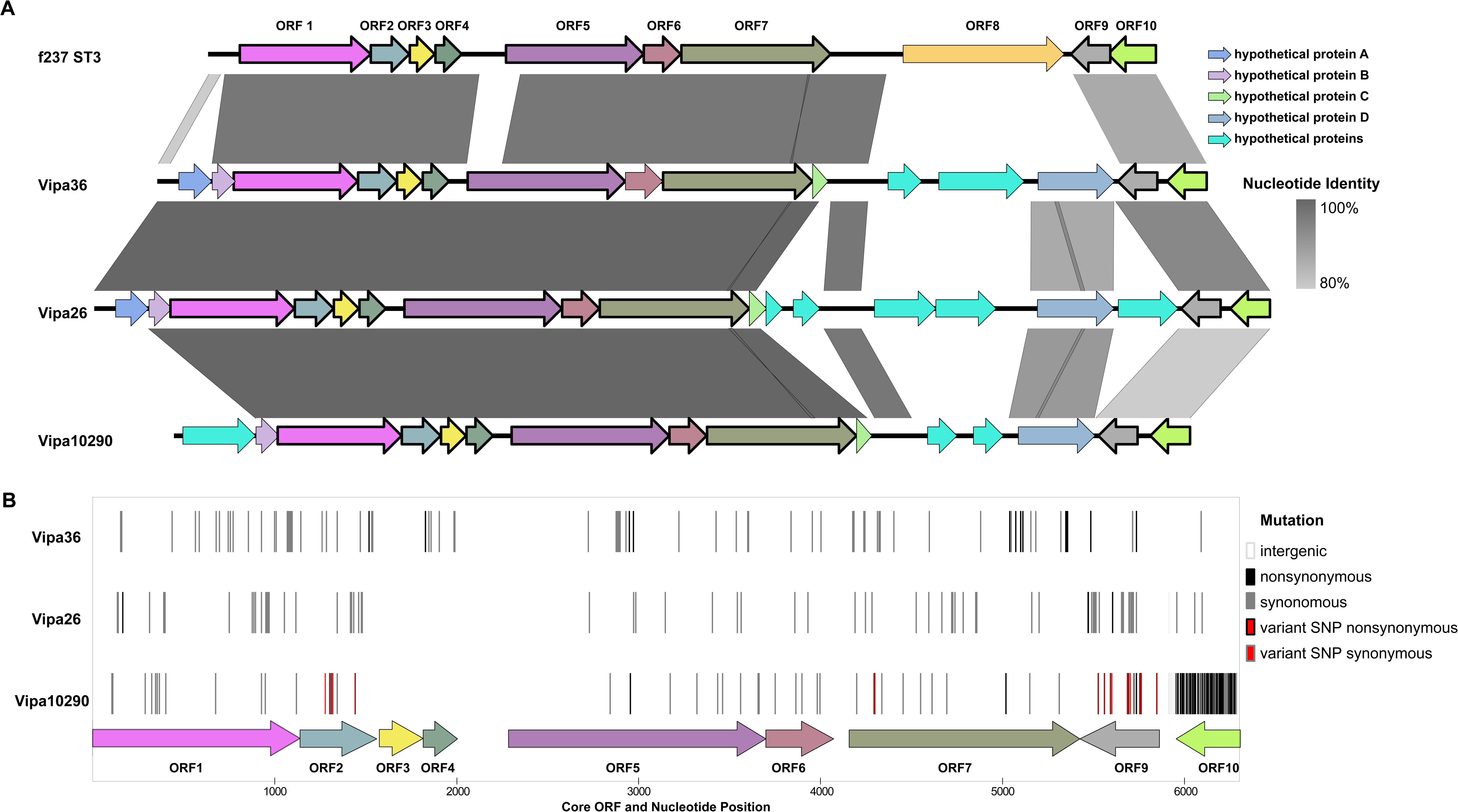
Comparisons of inoviruses in *V. parahaemolyticus* ST36. (A) Alignments of reference inovirus genomes depicting unique and shared content, and % nucleotide identity between Vipa36 (9,721 bp), Vipa26 (10,893 bp) and Vipa10290 (9,414 bp) as compared to f237 (8,784 bp). Orthologous ORFs are depicted by the same color except for non-homologous light blue hypothetical proteins. (B) Core inovirus genome (ORFs 1-7, and ORF9-10) single nucleotide polymorphisms (SNPs) were identified by comparison of quality ST36 genomes to the alignment consensus of reference inovirus genomes for Vipa36 (21 genomes), Vipa26 (24 genomes), and Vipa10290 (23 genomes) and used to assess spatial distribution of variation between the inoviruses and within each inovirus lineage. SNPs are mapped by location where ORF identities are labeled and colored to match those in (A), and SNP type designated by colored blocks (see Table S6). This indicates few non-synonymous mutations (see text for details) and uneven distribution of variation with ORF3, an intergenic region between ORF4 and ORF5, and the 5’ end of ORF5 being identical in all three phage. Only Vipa10290 displayed non-conserved core genome variation among ST36 prophage.

These inoviruses have a conserved central core (ORF 1-7) and ORF9 and ORF10 (Fig. 3A), and two variable regions. Whereas core phage gene functions were identifiable (Table 1), the functions encoded in variable regions were not discernable. Nucleotide variation in the inoviruses represented up to 46% of their host’s genome variation helping to explain differences between whole and core genome phylogenies (Fig. 1, S1 & S2) and population structures (Fig. 2 and Fig. S4). Though this content would be credited to recombination, surprisingly, genomes that lack inoviruses had a higher proportion of gene content assigned to blocks of recombination: 2.7% in 12310 and MAVP-1, and 2.0% in MAVP-23, compared to 1.3% in 10290, 1.4% in MAVP-26, and 1.8% in MAVP-36 (Fig. S1 and Table S3). Though divergent from one another, inovirus variation likely did not alter function: the 74 variant sites in the core of Vipa26 generated only three non-synonymous mutations, whereas the 78 variant sites in Vipa36 produced 13 non-synonymous mutations (Fig. 3B, Table S6). ORF1-7 are under purifying selection (codon-based Z test =9.425, p < 0.001) implying these are essential genes, and that their function is preserved. This also suggests the infections not cryptic, and the phage are likely still functional. Comparisons of core inovirus content from additional ST36 lysogens revealed 100% identity within the Vipa26 and Vipa36 lineages, whereas Vipa10290 had 18 non-conserved variable sites, perhaps reflecting its long history with a sizeable ST36 population (Fig. 3B).

**Table 1.**
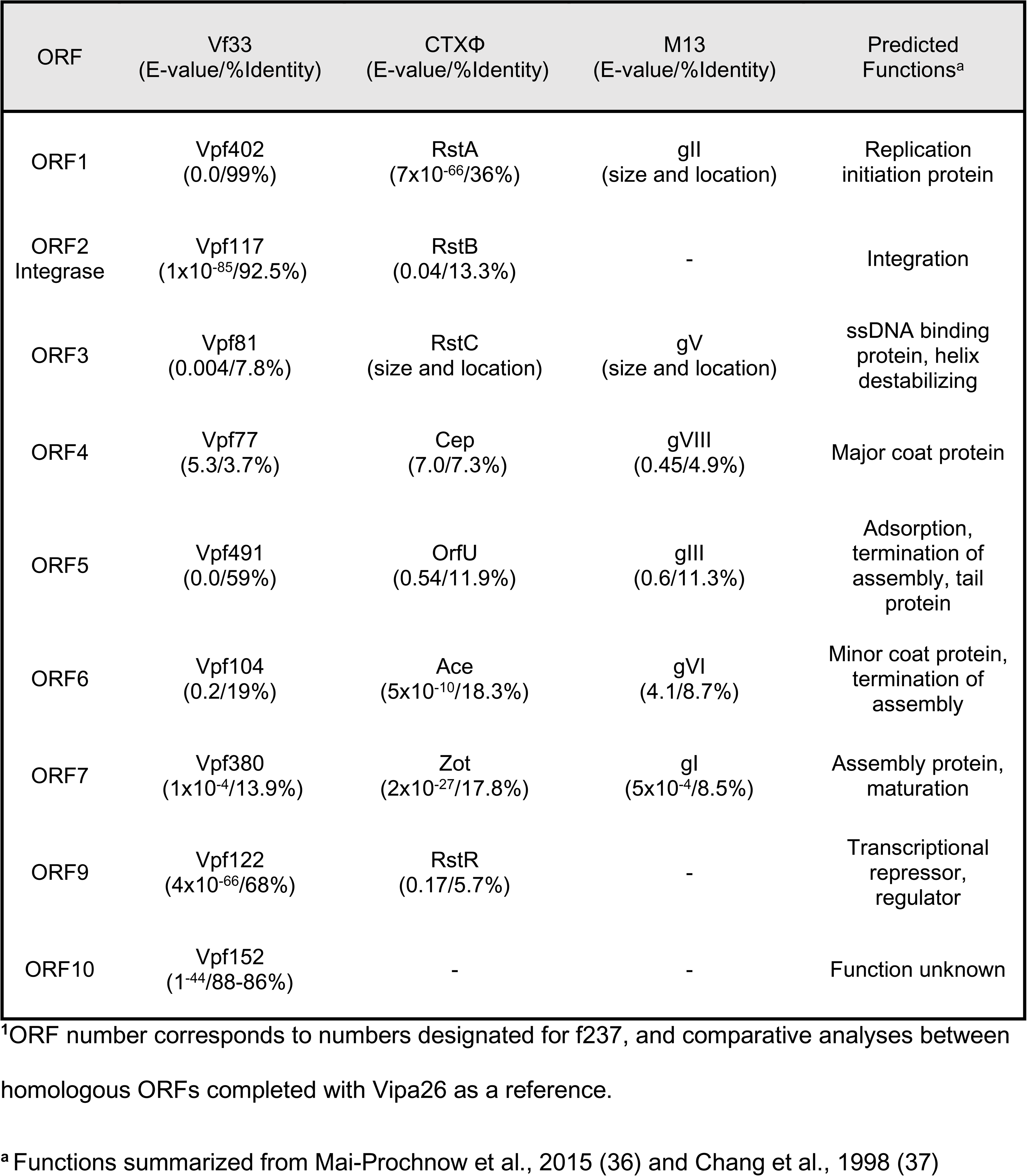
Inferred functions of inovirus genes ^1^.

| ORF | Vf33<br>(E-value/%Identity) | CTXΦ<br>(E-value/%Identity) | M13<br>(E-value/%Identity) | Predicted<br>Functions<sup>a</sup> |
| --- | --- | --- | --- | --- |
| ORF1 | Vpf402<br>(0.0/99%) | RstA<br>(7x10<sup>-66</sup>/36%) | gII<br>(size and location) | Replication<br>initiation protein |
| ORF2<br>Integrase | Vpf117<br>(1x10<sup>-85</sup>/92.5%) | RstB<br>(0.04/13.3%) | - | Integration |
| ORF3 | Vpf81<br>(0.004/7.8%) | RstC<br>(size and location) | gV<br>(size and location) | ssDNA binding<br>protein, helix<br>destabilizing |
| ORF4 | Vpf77<br>(5.3/3.7%) | Cep<br>(7.0/7.3%) | gVIII<br>(0.45/4.9%) | Major coat protein |
| ORF5 | Vpf491<br>(0.0/59%) | OrfU<br>(0.54/11.9%) | gIII<br>(0.6/11.3%) | Adsorption,<br>termination of<br>assembly, tail<br>protein |
| ORF6 | Vpf104<br>(0.2/19%) | Ace<br>(5x10<sup>-10</sup>/18.3%) | gVI<br>(4.1/8.7%) | Minor coat protein,<br>termination of<br>assembly |
| ORF7 | Vpf380<br>(1x10<sup>-4</sup>/13.9%) | Zot<br>(2x10<sup>-27</sup>/17.8%) | gI<br>(5x10<sup>-4</sup>/8.5%) | Assembly protein,<br>maturation |
| ORF9 | Vpf122<br>(4x10<sup>-66</sup>/68%) | RstR<br>(0.17/5.7%) | - | Transcriptional<br>repressor,<br>regulator |
| ORF10 | Vpf152<br>(1<sup>-44</sup>/88-86%) | - | - | Function unknown |

<sup>1</sup>ORF number corresponds to numbers designated for f237, and comparative analyses between homologous ORFs completed with Vipa26 as a reference.
<sup>a</sup> Functions summarized from Mai-Prochnow et al., 2015 (36) and Chang et al., 1998 (37)

With only one exception (A5Z905), all old-PNW clade members, including CTVP44C, contained Vipa10290 or related phage (Fig. 1, square). ST39, a member of the same clonal complex as ST36, also contains Vipa10290 whereas other clonal complex members (ST59 and ST21), share a distinct inovirus (Fig. S5, Tables 2, S9, & S10). Despite this history of lysogeny among the endemic population, no modern-day and definitively traced PNW ST36 isolates harbor inoviruses (Fig. 1 & Table S1 & S7) and this loss coincides with population diversification (Fig. 1). However, like GOM and SCC clade members, a total of nine other modern ST36 isolates collected between 2007-2016, some that dispersed to new locations and all others untraced, acquired eight other inoviruses (Fig. 1, Table 2).

**Table 2.** Summary of *Inoviridae* prophage in ST36 isolates.

| Strain | Inovirus name | NCBI Accession<br>(Genome coordinates) | Additional ST36<br>genomes with<br>inovirus | Other STs<br>with related<br>inoviruses |
| --- | --- | --- | --- | --- |
| 10290 | vB_Vipa10290 | AVOH01000008.1<br>(1370845..1378511) | Many | 38, 39 |
| MAVP-26 | vB_Vipa26 | MT188662 | Many | 1574, NF |
| MAVP-36 | vB_Vipa36 | MT188663 | Many | None |
| CDC_K5323G | vB_Vipa5323 | MIUF01000019.1<br>(188973..190181)<br>MIUF01000072.1<br>MIUF01000065.1<br>MIUF01000002.1<br>(4141..6171) | None | 752, 1132,<br>NF |
| CDC_K5308 | vB_Vipa5308 | MIUE01000114.1<br>(7376..8729)<br>MIUE01000031.1<br>(5201..8319) | None | 43 |
| 10-4255 | vB_Vipa71<sup>1</sup> | MT193890<br>NIXZ01000007.1<br>(181573..191867) | None | 43 |
| CDC_A8962 | vB_Vipa8962 | LHRO01000017.1<br>(70737..72090) | None | 327 |
| CDC_K5512 | vB_Vipa5512 | MIUZ01000070.1<br>(628..7658) | CDC_K5345G,<br>CDC_K5280 | 1713 |
| MEVP-10 | vB_Vipa10 | MT188666 | None | None |
| 1.146-15 | f237 | WSRX01000192.1,<br>WSRX01000239.1,<br>WSRX01000194.1,<br>WSRX01000150.1 | None | 3, 152, 809,<br>1473, NF |
| 1.220-16 | other | AAXNNA010000036.1 | None | None |

NF: Sequence type that is not yet assigned; None: no strains harboring phage.
<sup>1</sup>Named for ST43 strain MAVP-71

### The distribution of community-acquired phage offers clues of residency and dispersal

For the Atlantic ST36 populations, inovirus content was diagnostic of lineage and geography. Vipa26 was absent from only one member (MAVP-20) of the ST36 GOM clonal clade and present in only one distinctive isolate from a different location (CTVP25C) representing either basal or independent acquisition (Fig. 1 circles). Vipa36 (Fig. 1 triangles) is in all members of the SCC clade and only two isolates from this clade were traced to nearby locations. Because these inoviruses, like f237, are not widespread and have been maintained by these lineages tied to two locations, they are diagnostic of these ST36 subpopulations. In contrast, and like the modern PNW traced-ST36 isolates, no isolates of the LIS or MAC clonal clades harbor inoviruses.

The association of inoviruses with two Atlantic clades, and their absence in others, suggests that the founding subpopulation progenitors acquired phage upon Atlantic invasion only in some locations perhaps reflecting phage predation pressure. Environmental surveys revealed that although 22% of Northeast US *V. parahaemolyticus* isolates harbor inoviruses, this proportion varied by location (50). Proportions were lower in LIS (10%), than in GOM and SCC (30% and 42% respectively). Whereas no environmental isolates other than ST36 harbor Vipa36, multiple non-ST36 isolates collected in NH as early as 2008 contain Vipa26 (Table S1, Fig. 4, & S6). This suggests Vipa26 was acquired locally as supported by its absence from VpG-1 (Fig. 1, Table S1 & S7).

**Figure 4.**
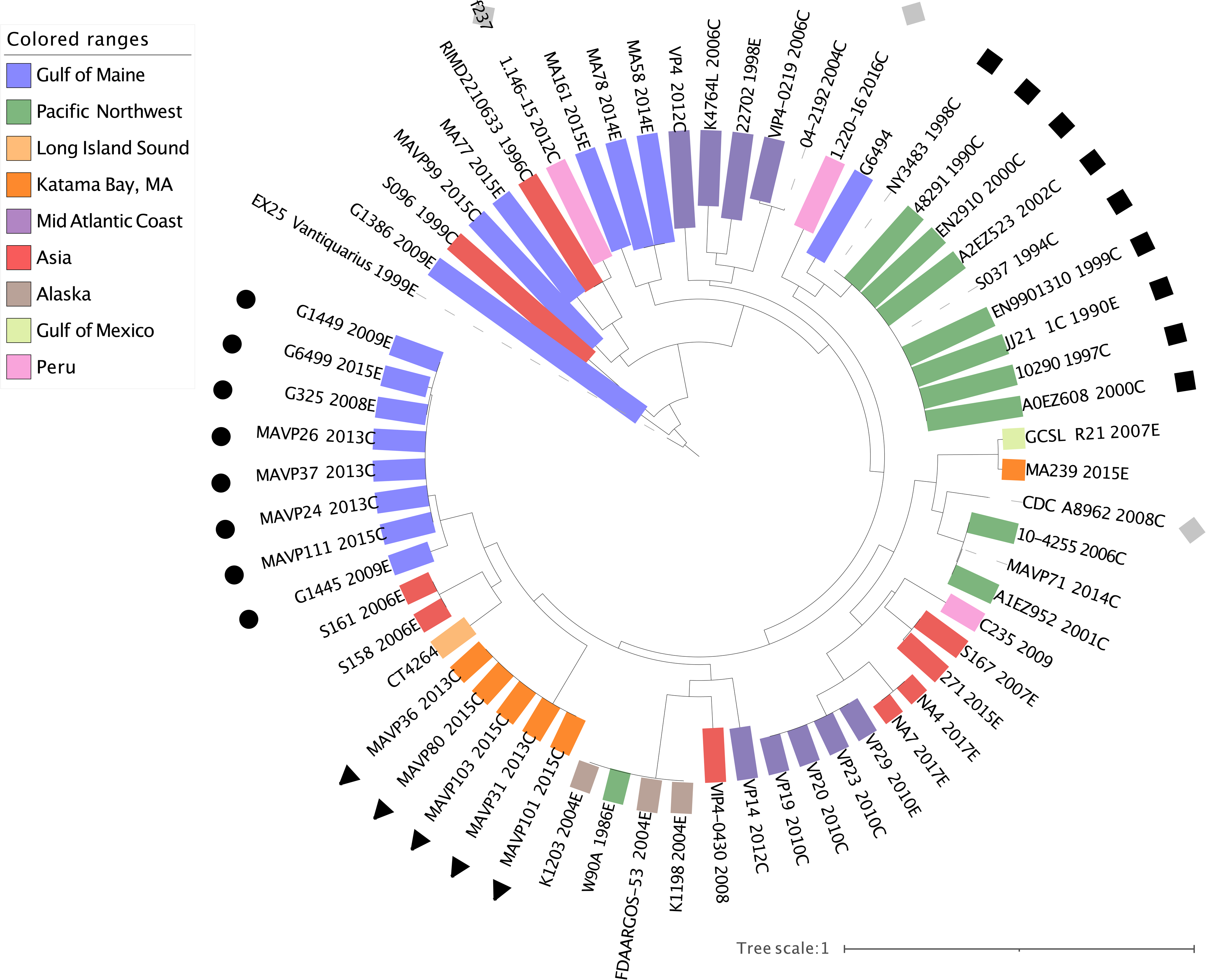
Whole genome phylogenies of representative inoviruses from diverse locations and sequence types. Whole genome phylogenies of select inoviruses from environmental (E) and clinical (C) isolates of *V. parahaemolyticus* labeled by name and year of isolation. Bootstraps criterion met after 350 replicates. Environmental traceback color coding and symbols, representing named phage corresponds to those in Figure 1 where circle is Vipa26, square is Vipa10290, triangle is Vipa36 and grey square is other (Vipa8296). ST36 isolates are in bold.

Because comparative phage phylogeny could elucidate the history of ST36 as members moved through other populations, we expanded our search for inoviruses among available genomes of *V. parahaemolyticus.* These prophage genomes added to the *Inoviridae* diversity (Fig. & S5) but display little discernable phylogeographic population structure in that related phage are found in both the Atlantic and Pacific *V. parahaemolyticus* populations, and phage in Asian isolates group among most branches, with the exception of Vipa10290. Two ST36 isolates harbor two different inoviruses maintained by members of the PNW co-resident but unrelated ST43 (Table 2, Fig. S5). Though most members of the two ST36 lineages reported in Peru also lack inoviruses, including a 2011 environmental isolate related to the Spain lineage (45), one clinical isolate of this lineage harbors a Vipa10290-related inovirus, whereas a second clinical isolate (1.146-15), harbors f237 (Fig. 4).

The phylogenomic distribution of inoviruses in all publicly available *V. parahaemolyticus* genomes indicates they are common, with 46% of genomes containing one or more inovirus (Fig. 5, Fig. S6, and Table S10) yet distributed unevenly. Many lineages harbor a persisting inovirus (Fig. 5 and Fig. S6). A few lineages harbor two evolutionary distinct inoviruses concurrently suggesting that protection from superinfection, a common attribute of both temperate phage and inoviruses (51, 52), is not absolute (Fig. S5). In addition to Vipa26, a second inovirus is present in multiple, unrelated STs (Fig. S6). Even though most STs have an inovirus prophage that they propagate vertically, the inoviruses of some isolates were replaced by another, like ST36 (Table 3, Fig. S6). Notable among these are emergent pathogenic lineages including ST43 and the prevalent Atlantic endemic ST631 lineage that has caused illnesses along the North American Atlantic coast (4), the pandemic complex (ST3) and a diverse and far-spreading Asian lineage that is now an Atlantic resident (ST8).

**Figure 5.**
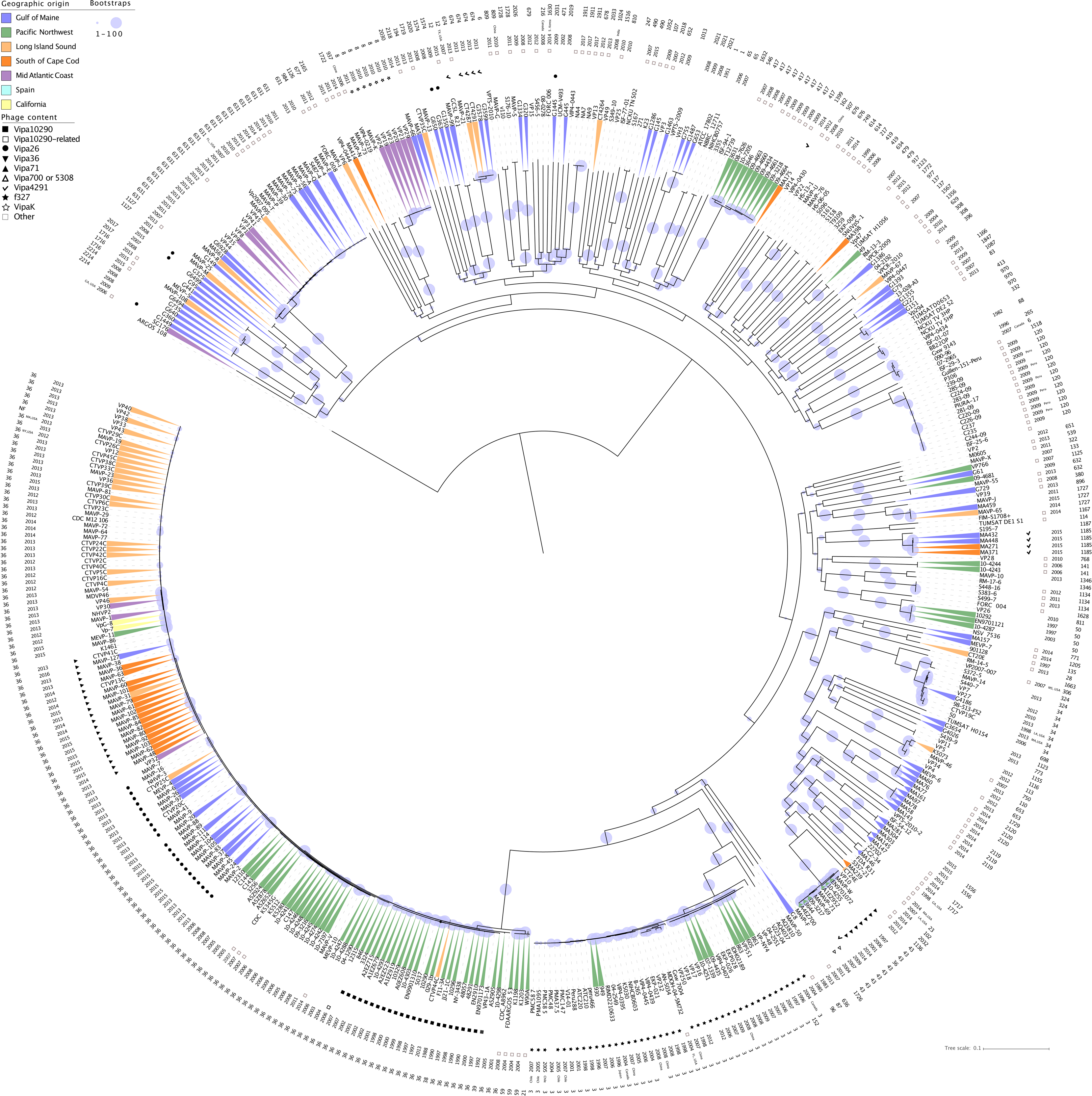
Maximum-likelihood tree of diverse *V. parahaemolyticus* isolates with their inovirus content. A maximum-likelihood (ML) phylogeny was built on 1,025,281 genome SNPs where 250 bootstraps (criterion met) were mapped onto the best scoring ML tree. Isolates are colored to correspond to geographic region as in Fig. 1, where no color indicates unknown or ambiguous origin. Symbols next to strains indicate unique inovirus content. Outermost label indicates the sequence type (ST) where NF or unlabled = sequence type not found or not known, followed by year of isolation, if known.

**Table 3.** Lineages other than ST36 with diverse phage genotypes.

| Sequence type | Number of unique Inovirus prophage in lineage | Sequenced strains without prophage |
| --- | --- | --- |
| 631 | 4 | Yes |
| 674 | 4 | No |
| 43 | 5 | Yes |
| 12 | 4 | No |
| 3 | 2 | Yes |
| 8 | 2 | Yes |
| 308 | 2 | Yes |
| 120 | 2 | Yes |

## Discussion

Here we demonstrate that changes in inovirus prophage accompanied diversification, competitive replacement, dispersal, and Atlantic invasion of *V. parahaemolyticus* ST36. This suggests that inoviruses could countribute to the emergence of new lineages with the potential to spread more broadly. The misleading nature of disease reporting that does not align illnesses with environmental reservoirs and declining availability of recent ST36 genomes from the US PNW have been obstacles to deciphering the ecological context of ST36 evolution and spread. We addressed these limitations using genomes from isolates reported in Canada and other US states that were publicly available and by augmenting available genomes with a collection of ST36 clinical isolates from the Northeast US. We carefully curated environmental traceback to advance our understanding of ST36 ecological expansion into the Atlantic Ocean.

Our analysis indicates five or more unique ST36 progenitors diversified in the Pacific and then translocated into the Atlantic to found subpopulations (Fig. 1, S2, 2, and S4). The earliest sequenced ST36 isolate traced to the Atlantic in 2006 (VpG-1) and several PNW strains share a similar pattern of mixed ancestry with this isolate and group with the GOM clade suggesting its progenitor may have arrived prior to 2006 with little effect on epidemiology until after the anomalously mild winter of 2011-2012 (1, 2). Isolates traced to SCC, LIS and MAC are more related to each other, and multiple Pacific-traced isolates group within or adjacent to their clades (Fig. 1, 2, & S2). This pattern is most simply explained by divergence of these populations’ ancestors in the Pacific prior to introduction into the Atlantic (Fig. 1, 2 & S2). Clade V’s shared ancestry with CA and PNW isolates further implicates the PNW as the source population. And although imperfect trace back or mislabeled genomes could explain some unexpected patterns, nearly every Pacific-traced relative of Atlantic residential populations were isolated, reported, and sequenced independently and prior to Atlantic isolation (Table S1). Although our interpretations of population evolution and dispersal contrast with previous reports (9), this is mostly explained by differences in geographic assignment of isolates to environmental sources instead of the readily available location of disease reporting as our goal was to understand dynamics affecting reservoirs of emergent pathogenic lineages (See details of Table S1). Traceback confirmation from CA (Table S1), disease reporting in Mexico (53) and environmental isolation in Peru (9, 45) further corroborate that these lineages diversified in the Pacific. And though missing intermediate populations could suggest unknown reservoirs exist, it is more likely that sampling bias, a decrease in availability of PNW genomes, decreased clinical prevalence of ST36 in the PNW, and lack of surveillance there have compounded challenges with deciphering this pathogen’s evolution and spread. Based on mounting evidence that most if not all lineages arose in the PNW, this warrants further examination of the PNW environmental reservoir to manage further spread.

Even as increasing ocean temperatures and weather events have been tied to both seasonal illnesses by *V. parahaemolyticus* and its spread to new locations (9, 54–57), the stable and non-mixing distribution of Atlantic ST36 lineages suggests that environmental fitness that promotes persistence may be their key to success. It is notable that in BC Canada where many diversified ST36 isolates were first reported, rising infection rates were not fully explained by temperature models but rather attributed to new pathogenic strains (58, 59). Like the Northeast US, the MAC experienced a warming trend in 2012 (9), nonetheless this location has not experienced persistent problems with ST36 (1, 10). Combined, we believe warming ocean temperatures are only part of the picture and that underexplored ecological interactions are at play in driving *V. parahaemolyticus* invasion and persistence (60). Even in locations in the Atlantic with recurrent illness, environmental prevalence of ST36 is extremely rare (Table S1) illustrating how low-level persisting populations can change epidemiology. Identification of unique ecological and population contexts of the MAC compared to more northern locations could provide useful insight.

Phage loss and later sequential phage replacement is one of the most striking and common features of ST36 lineage succession in the PNW and population expansion into the Atlantic. Nearly all old PNW lineage members harbor Vipa10290 (Fig. 1 and Table S1), whereas strains of the new ST36 lineages isolated in the PNW after 2002 lack inoviruses, though other phage are not uncommon (Table S3). The presence of Vipa10290 in other clonal complex members could indicate a basally acquired phage was lost during lineage diversification. Though inovirus excision is infrequently reported (61, 62), laboratory passage of ST36 can lead to both phage curing and loss of protection from superinfection (63). The lack of inovirus-harboring PNW ST36 population members indicates they ultimately were at a disadvantage in the context of a changing population even though other residents concurrently maintained inoviruses (e.g., ST43, Table 2) and potentially transmitted these to a few ST36 members. The subsequent acquisition of inoviruses by diverging ST36 lineages (Table 2) indicates ST36 remained susceptible to infection, though the persisting inovirus-free state of modern-PNW ST36 lineages suggests that resistance may have evolved during their diversification, as has been described in other sympatric *Vibrio* populations (64). In contrast, that the progenitors of two clinically prevailing Atlantic ST36 lineages acquired inoviruses prior to clonal expansion, while ST36 populations in the LIS where inoviruses are less abundant did not, indicates phage acquisition was not requisite for success, perhaps reflecting differences in phage predation and other population dynamics. Even so, whereas resident ST36 from SCC and GOM have continued to cause sporadic disease, infections from LIS have precipitously declined, in part reflective of successful management.

Inovirus prophage confer obvious costs through persistent non-lethal virion production (61), suggesting their loss would be favored; and yet inoviruses have been maintained by prevalent lineages for decades (Table 2 & 4, Fig. 5). This could reflect their fastidious nature, but the success of the two northernmost Atlantic populations harboring inoviruses contrasting with the unsuccessful lineages that lacked them signals alternatives are possible: that inoviruses may confer advantages in some as yet undefined context. For example, shed inovirus virions contribute to biofilm matrix (65) thereby promoting virulence and antimicrobial resistance, but also potentially enhancing survival during long-distance movement on particles (52, 54, 66–68). Shed inoviruses can mislead human immune responses and decrease host ability to clear bacterial infections, thereby enhancing virulence (69). Furthermore, variable content in some inoviruses is linked to anti-predation (17, 70) and virulence (22, 27, 39). We find it intriguing that succession in the PNW and loss of inoviruses by ST36 accompanied its clinical decline there. Future functional studies with these inoviruses are essential to discern whether any of these possibilities are at play.

Prophage can also modify their hosts in ways that alter their capacity for horizontal gene transfer (HGT). Prophage can protect against superinfection, including by unrelated lytic phages (27, 61, 71) thereby blocking transduction. They also modulate, and concurrently block conjugative pili (72). The concept that loss of prophage-conferred immunity contributed to ST36 evolution is attractive considering DNA release by predation of non-lysogen susceptible hosts could promote transformation through natural competence, though the conditions of natural competence by *V. parahaemolyticus* are unknown (73–75). Type IV pili which are the receptor for some inoviruses also facilitate transformation mediated HGT (51, 52). If the newly inovirus-less ST36 lineages were more vulnerable to phage infections, phage transduction, and conjugal transfer alike this could have contributed to its rapid diversification (Fig. 1) whereas translocating lineages would likely not have evolved mechanisms to resist resident Atlantic phage facilitating re-acquisition (31, 52). In keeping with this premise, many of the most highly divergent ST36 individuals lacking inoviruses evolved by multiple recombination events including an isolate of the MAC clade (VP30). Accumulation of non-inovirus prophage was notable in the early diverging Spain lineage (clade I) (9). And whereas the inovirus-harboring SCC and the GOM clades remain remarkably stable, as do their phage, Atlantic isolates lacking inoviruses, including the single GOM clade member (MAVP-20) and many members of the LIS population exhibit complex admixture (Fig. S4). In contrast, the inovirus-harboring old-PNW relic CTVP44C isolated in LIS in 2013 is virtually unchanged from the last known members of this lineage from the PNW that were isolated in 2002 (Fig. 1, Table S1). Furthermore, a 2005 isolate of this same lineage lacking Vipa10290, A5Z905, is among the most highly divergent ST36 strains (Fig. 1, 2, S2). Similarly, pandemic O3:K6 strains, which stably maintain f237, reported in the Americas between 1996-2012 display remarkably little genetic variation from the type strain for this lineage RIMD 2210366 (76). Though these data only establish correlation of phage absence and diversification, it alludes to the possibility that inovirus prophage that provide some level of resistance to superinfection and block conjugation could thwart HGT, and by contrast, loss of inoviruses could remove barriers to recombination and promote rapid diversification. The selective conditions that favor inovirus acquisition and maintenance or loss would thereby influence some mechanisms that could promote rapid evolution, potentially of more or even less pathogenic potential.

These analyses suggest an intriguing connection of inovirus prophage to pathogen evolutionary dynamics, alluding to the possibility that inoviruses could be gatekeepers of HGT in natural ecosystems driving changes in human disease epidemiology. With broader population analysis and functional studies, this could lay the foundation for a new evolutionary paradigm where inovirus prophage-mediated immunity is a major governing force beyond the appreciated roles of phage in selective predation (20–22) and phage conversion (23–27). The apparent ability of ST36 and some other lineages (Table 2 & 3) to transition between phage infection states or replace their phage could generate flexibility in the balancing of fitness tradeoffs under different selective regimes that are not yet understood (29, 64). The high prevalence of inoviruses not only among the global population of *V. parahaemolyticus* (Fig. 5) but universally in other bacteria (27) with little clarity of how inoviruses impact the ecology of their hosts calls for more mechanistic analyses which could reveal varied roles for inoviruses in promoting evolution and competitive dominance of emerging pathogens.

## Materials and Methods

### Vibrio parahaemolyticus genomes

Regional clinical strains spanning 2010-2018, and trace back were acquired from public health laboratories (2). Only isolates with single source traces were assigned a location. Environmental isolates were collected between 2007-2015 (1, 2). Isolates were identified as *V. parahaemolyticus* by *tlh* gene amplification (77, 78) and identity confirmed by genome sequencing. Publicly available genomes were acquired from NCBI (https://www.ncbi.nlm.nih.gov/genome/681; August 2019).

DNA was extracted using the Wizard Genomic DNA purification Kit (Promega, Madison WI USA) or by organic extraction (2). Sequencing libraries were prepared as described (79). Genomic DNA was sequenced using an Illumina – HiSeq2500 device at the Hubbard Center for Genome Studies at the University of New Hampshire, using a 150bp paired-end reads with de-multiplexing and quality filtering prior to analysis. The *de novo* genome assembly were performed using the A5 pipeline (80), and annotations assigned with Prokka1.9 using “*Vibrio”* for the reference database (81). The sequence types were determined using the SRST2 pipeline (82) or using assemblies (83) referencing https://pubmlst.org/vparahaemolyticus/ (84). Reference inoviruses were extracted from genomes sequenced using the Pacific Biosciences RSII technology and with Illumina short read error correction as described (4).

### V. parahaemolyticus Phylogenetic Relationships

Because all ST36 strains are closely related (clonal) and recombination contributed to the recent lineage divergence, ST36 strain relationships (Fig. 1) and all *V. parahaemolyticus* relationships (Fig. 5) were determined using reference-free, whole genome alignments using kSNP 3.1 (85) and maximum likelihood phylogenetic trees built by RAxML (86) and visualized with iTOL(87). The optional Kchooser script provided in kSNP3.1 determined the optional kmer size of 19, for each alignment dataset. Whole genome trees were rooted with *Vibrio alginolyticus* ARGOS_108 (hidden after rooting in iTOL). Maximum likelihood trees were inferred using the GTR-GAMMA model of nucleotide substitution, and the convergence criteria autoMRE option for automatically determining a sufficient number of rapid bootstrap replicates. RAxML completed a thorough ML search after optimization on every 5^th^ bootstrapped tree, which were mapped on the ML tree with the best likelihood (“raxmlHPC-PTHREADS-AVX -m GTRGAMMA -f a -N autoMRE -x 12345 -p 12345” options). Gene/features were visualized with EasyFig 2.2.0 (88, 89).

In parallel, homoplastic regions resulting from recombination were identified in genomes by alignment to reference strain 10296 (90) (summary table S2) using Gubbins 2.3.4 (91) and removed, and the relationships of strains determined based on nucleotide SNP alignments generated with Gubbins 2.3.4 (91) (Fig. 2). Phylogenies were inferred as above, except using the GTR-CAT nucleotide substitution model (because the alpha parameter was >10). ML search and optimization, rapid bootstrap criterion, and mapping of bootstraps were done as described above. Unmapped reads were assembled using SPAdes 3.13.1 (92) and annotated using Prokka 1.14(81) (Table S4).

Population structure of ST36 was performed using whole and core SNPs genetic matrixes generated from SNP alignments in the R statistical package LEA (49, 93, 94) (Fig. 2). To determine the number of genetic clusters best explained by the distribution of genomic variation, we explored ancestral populations (*K*) of between 1-20 and used the entropy criterion to evaluate the quality of fit of the statistical model to the data and selected the probable number of populations based on minimal entropy. Once strains were assigned to populations, the least-squares estimates of ancestry proportions were plotted using ggplot2 (95).

### Inovirus Identification

ST36 strains were compared to reference 10290 (5) using Breseq (96). Unique prophage were classified as belonging to the family *Inoviridae* proposed subfamily *Protoinoviridae* (27) and assigned unique names (97, 98) (Table 2). All *V. parahaemolyticus* genomes were independently searched against a BLASTn (99) database constructed using the central core (ORF1 – ORF7) of vB Vipa-26. Nucleotide sequences of 745 phage were extracted and to determine if they were unique, maximum likelihood phylogenies using the GTR-GAMMA model for substitution and rapid bootstraps were constructed via RAxML on nucleotide SNPs generated by kSNP (Fig. S6). Rapid bootstraps reached criterion after 350 replicates and were mapped on the tree as described above. The complete prophage assemblies of unique ST36 inoviruses were named from the highest quality assembly as were other prevalent inoviruses (Table S9). Relatedness of select whole phage genomes was visualized using the Genome-BLAST Distance Phylogeny (GBDP) method (100) to conduct pairwise comparisons of the nucleotide sequences under settings recommended for prokaryotic viruses (101). Trees were rooted at the midpoint (28) and visualized using iTOL (87) (Fig. S5).

To determine if phage evolved under selection, a Nei-Gojobori codon-based Z test (102) was performed in MEGA 6 (103). Protein sequences for the phages Vf33 (NC_005948.1) (104, 105), CTXΦ (MF155889.1)(27), and the type strain for *Inoviridae* M13, which infects *Escherichia coli* (GCF_000845205.1) were compared using BLASTp (99).

Oligonucleotide primers were designed for multiplex amplification with the species-specific *tlh* primers (77, 78) where primers annealing within ORF3 within ORF5 identified inovirus presence, and primers annealing within HypD and ORF9 produced different size amplicons diagnostic of Vipa26 and Vipa36 (Table 4). Isolates were screened with 0.2µM primer annealed at 55°C with 1.5 min extension and phage identity confirmed by genome sequencing.

**Table 4.**
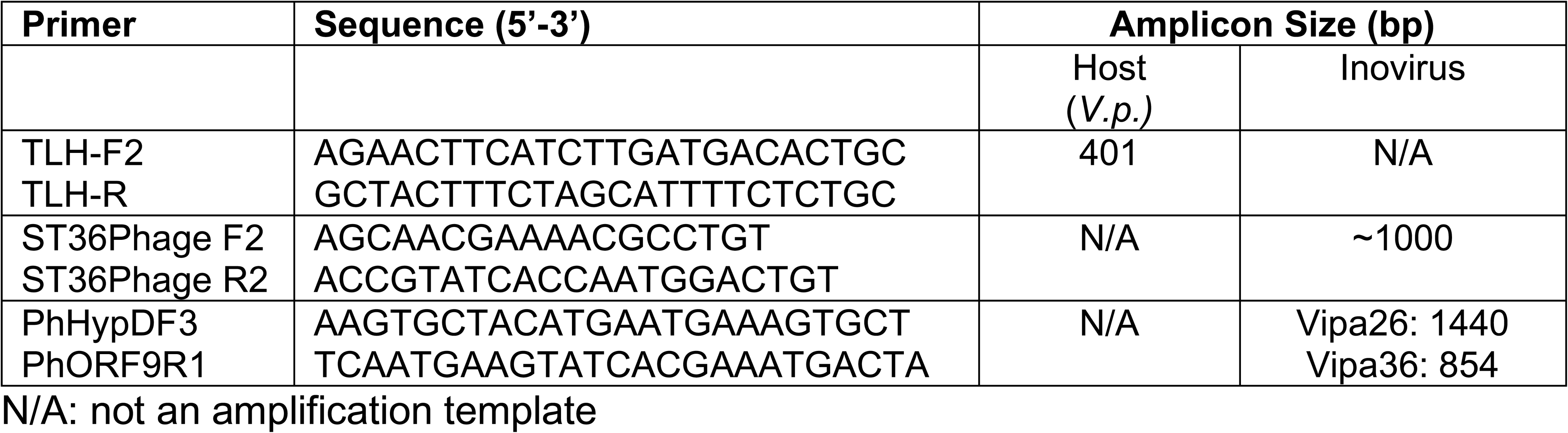
Multiplex PCR for detection of inoviruses.

### Data Availability

Genome accessions are listed in Tables S1, S7, S8 and Table 2.

## Acknowledgments

Partial funding was provided by the National Sea Grant College Program and projects R/SFA-4, R-HCE-3, and R/SFA-13, and by the New Hampshire Agricultural Experimentation Station. This work (scientific contribution number 2995) was supported by the USDA National Institute of Food and Agriculture Hatch projects NH00609 [accession number 233555], NH00625 [accession number 1004199], NH00658 [accession number 1013479], and NH00698 [accession number 1023165]. We thank Narjol Gonzalez-Escalona for helpful insight and feedback; Kanwit Kohl, Stephen Combes and Heather Grieser from the Maine Center for Disease Control and Prevention for strains and information; Tracy Stiles, Eric Hickey, Coleen Murphy, Michael Moore, and Jana Ferguson from the MA Department of Public Health for helpful insight, strains and epidemiological information; Jeff Kennedy and Christian Pepitas from the MA Division of Marine Fisheries for insight and guidance; Christopher Nash and Robert Atwood from the New Hampshire Department of Environmental Services and the Fish and Game Department, and Colleen Smith from the New Hampshire Department of Health and Human Services for information and discussion. We also thank Zoe Pavlik for curation of genome accessions.

Role of authors: JM, AM, RF, FX, CS, KD, JH, VC, SJ, CW collected samples, performed and interpreted analyses, curated databases, and constructed visuals. JM and AM drafted the manuscript and CW and RF completed writing the manuscript. SJ, VC, and CW designed and directed the studies. All coauthors reviewed and edited the manuscript for accuracy and completeness.

## Notes

### Competing Interest Statement

The authors have declared no competing interest.

### Summary of Updates

Fig. 3 has been updated, and details of phylogenetic construction added.

