## Supplemental Figures for "*Inoviridae* prophage and bacterial host dynamics during diversification, succession and Atlantic invasion of Pacific-native *Vibrio parahaemolyticus*"

### Figure S1

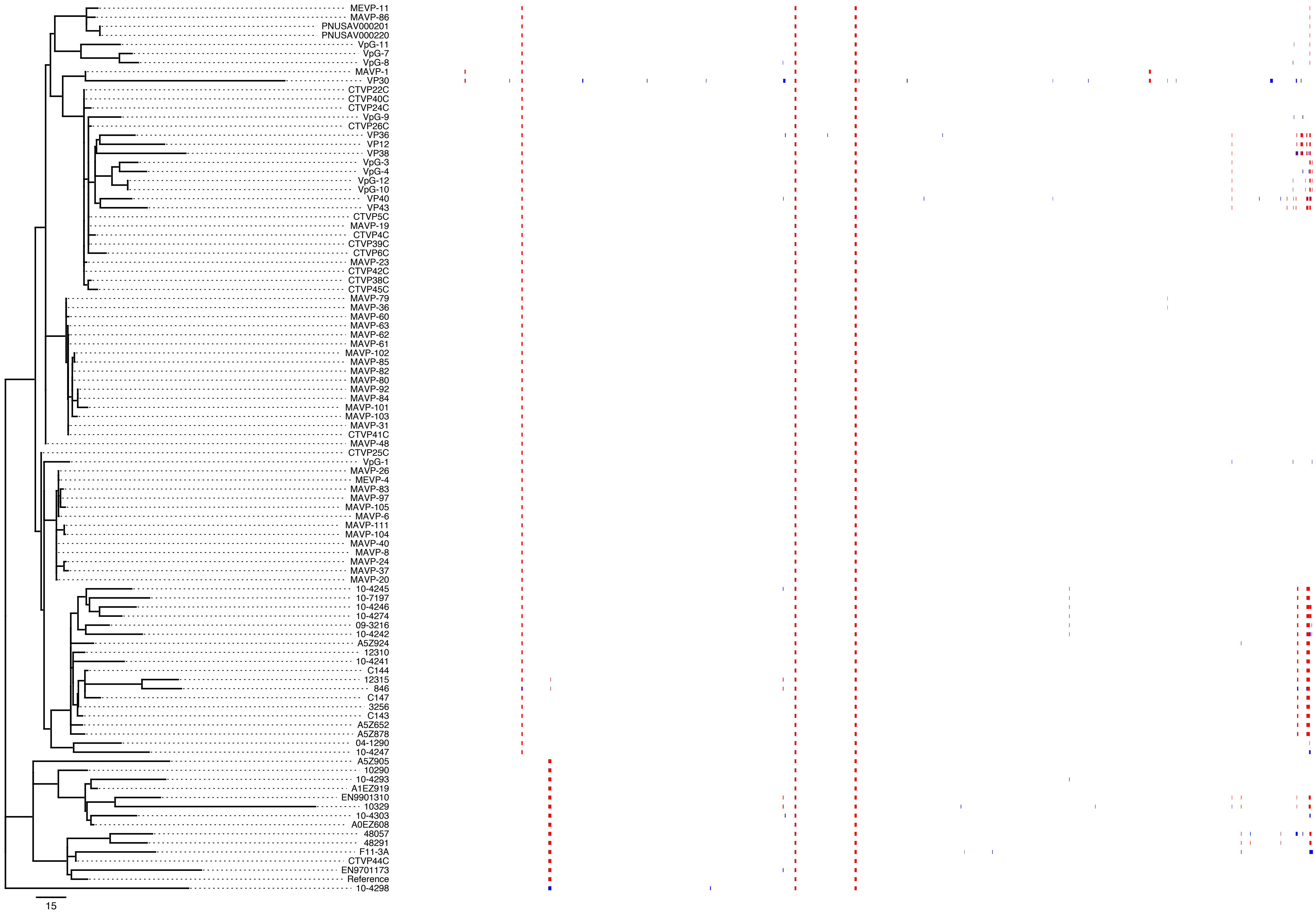


**Supplemental Figure S1. Phylogeny of ST36 excluding regions of recombination.** A maximum-likelihood (ML) phylogeny was built on SNPs identified in non-recombining regions (non-colored regions) and excluding regions of recombination exhibiting a higher SNP density (colored blocks) as identified by Gubbins (Croucher et al., 2015). Red blocks indicate recombination within a clade of related isolates, whereas blue blocks indicate recombination with isolates that were absent from the analysis

### Figure S2

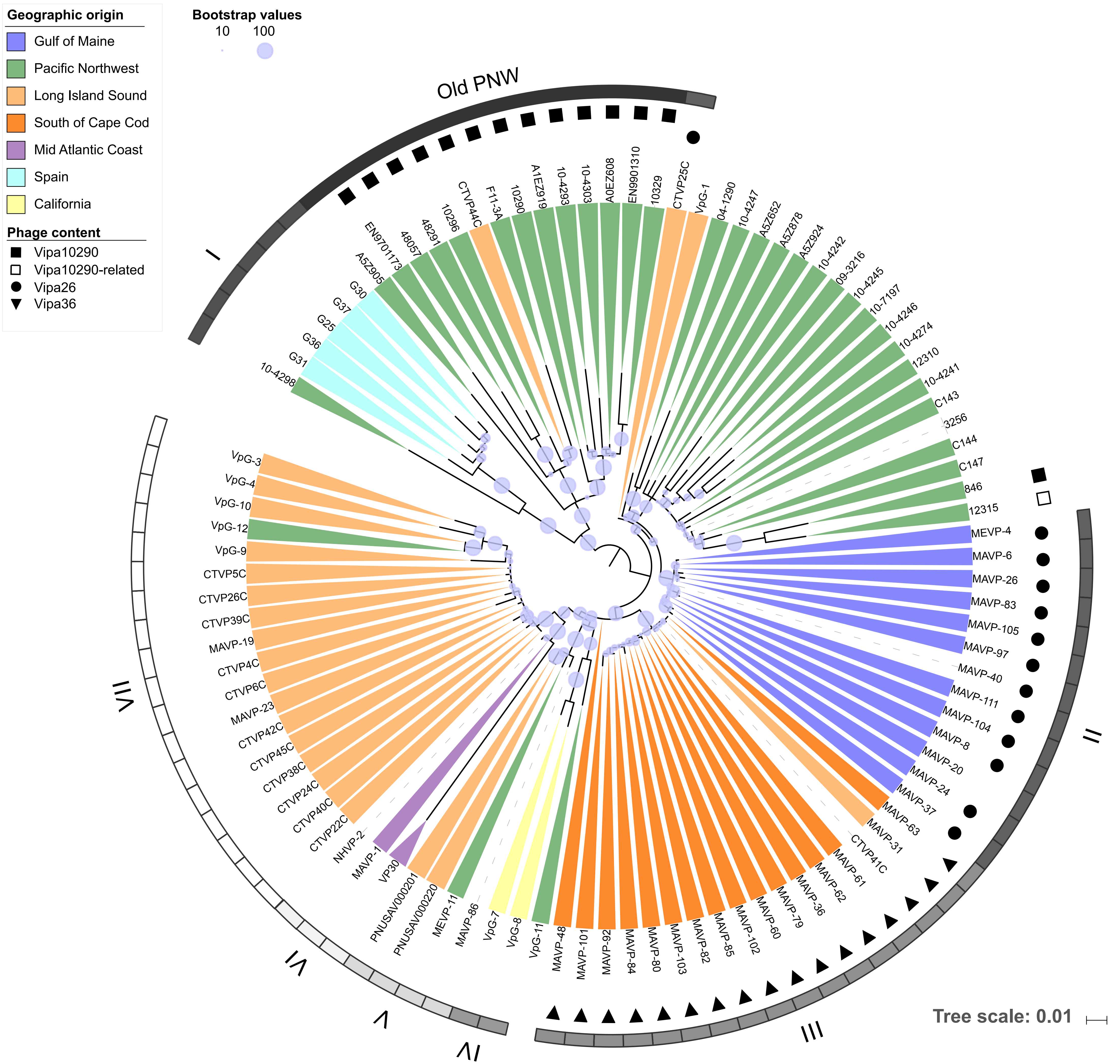


**Supplemental Figure S2 Maximum-likelihood phylogeny of ST36 *Vibrio parahaemolyticus* core non-combining genomes** A maximum-likelihood (ML) phylogeny was built on SNPs identified in core non-recombining regions identified by Gubbins. 1000 bootstraps were mapped onto the best scoring ML tree. Isolates are colored by geographic region, where available (Table S1) where no color indicates unknown or ambiguous origin. Symbols next to strains indicate unique Inovirus content. The ancestral PNW ST36 population is identified by a black bar, whereas seven different lineages associated with translocation events (I – VII) are identified by greyscale bars. Though LIS clade isolates sequenced in different years as part of this study had short terminal branches, genomes from strains traced to the same location (CT) and year (e.g. VpG-10, VP38), reported in other states and whose genome sequences were publicly available often exhibited very long branches suggestive of artifactual variation that may corrupt analyses, justifying removal of these strains from further analysis.

**Figure S3**

**A**

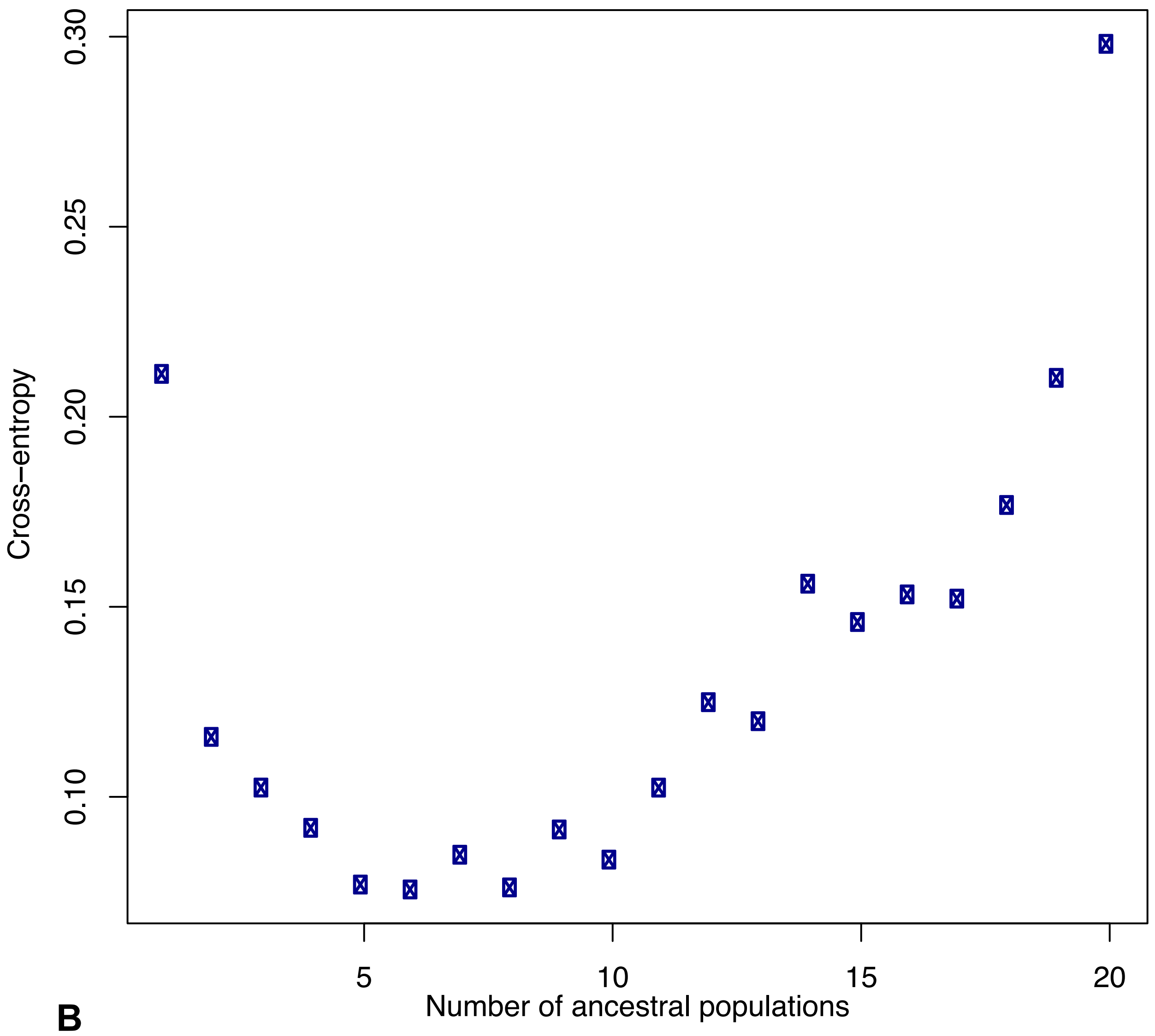


**B**

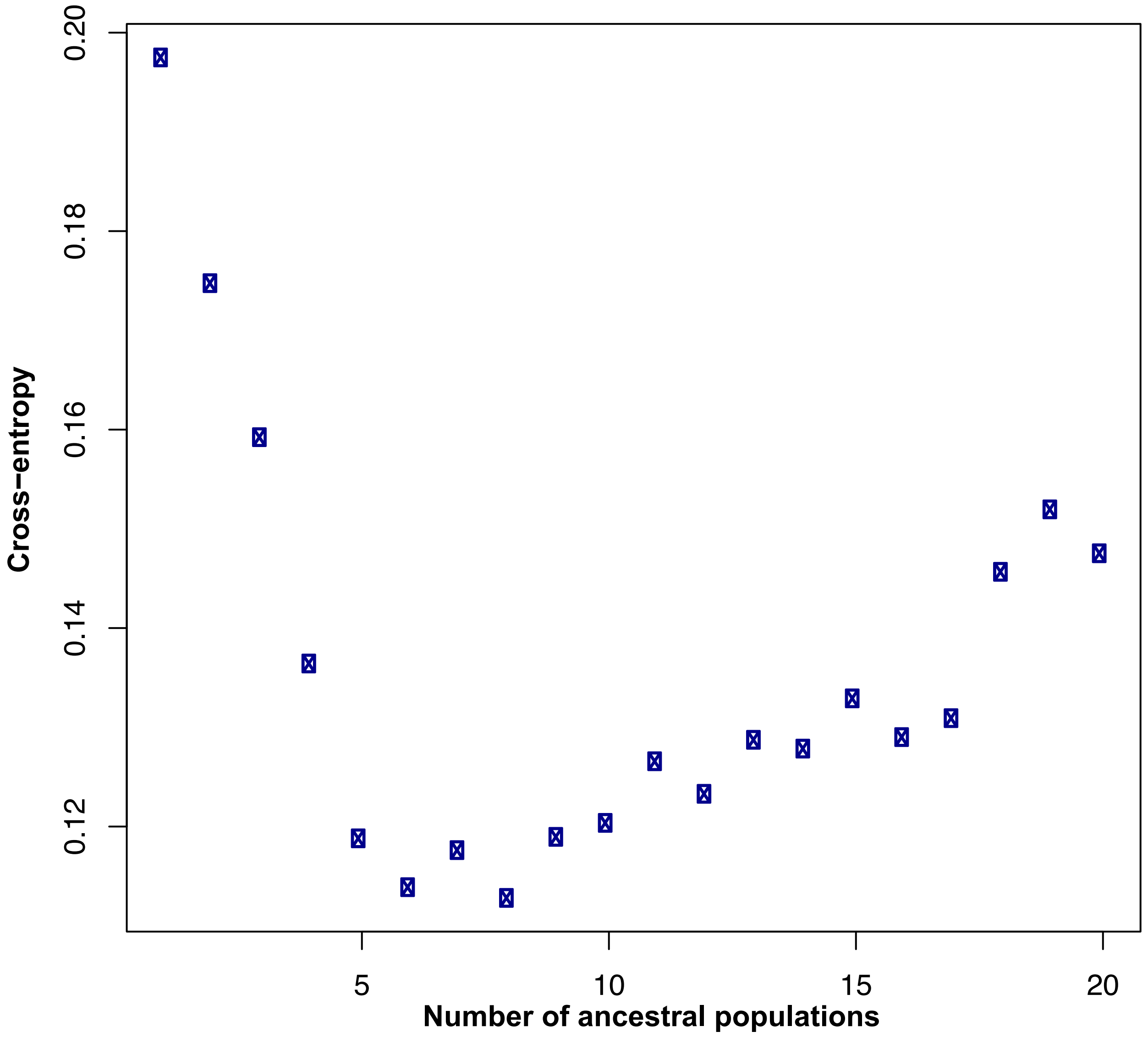


**Supplemental Figure S3. Cross-entropy derived from the coancestry distribution for an assigned number of ancestors that contributed nucleotide variation in the ST36 population**. Cross-entropy was determined by LEA (Frichot and François, 2015) by empirical evaluation of the likelihood that between 1-20 ancestors explains the distribution of core, non-recombining (A) or whole (B) genome variation among the global population of ST36. The lowest entropy was observed at 6 and 8 (Fig. 2) ancestral populations for core, non-recombining data, and 8 ancestral populations (see Fig. S4) for whole genome data.

### Figure S4

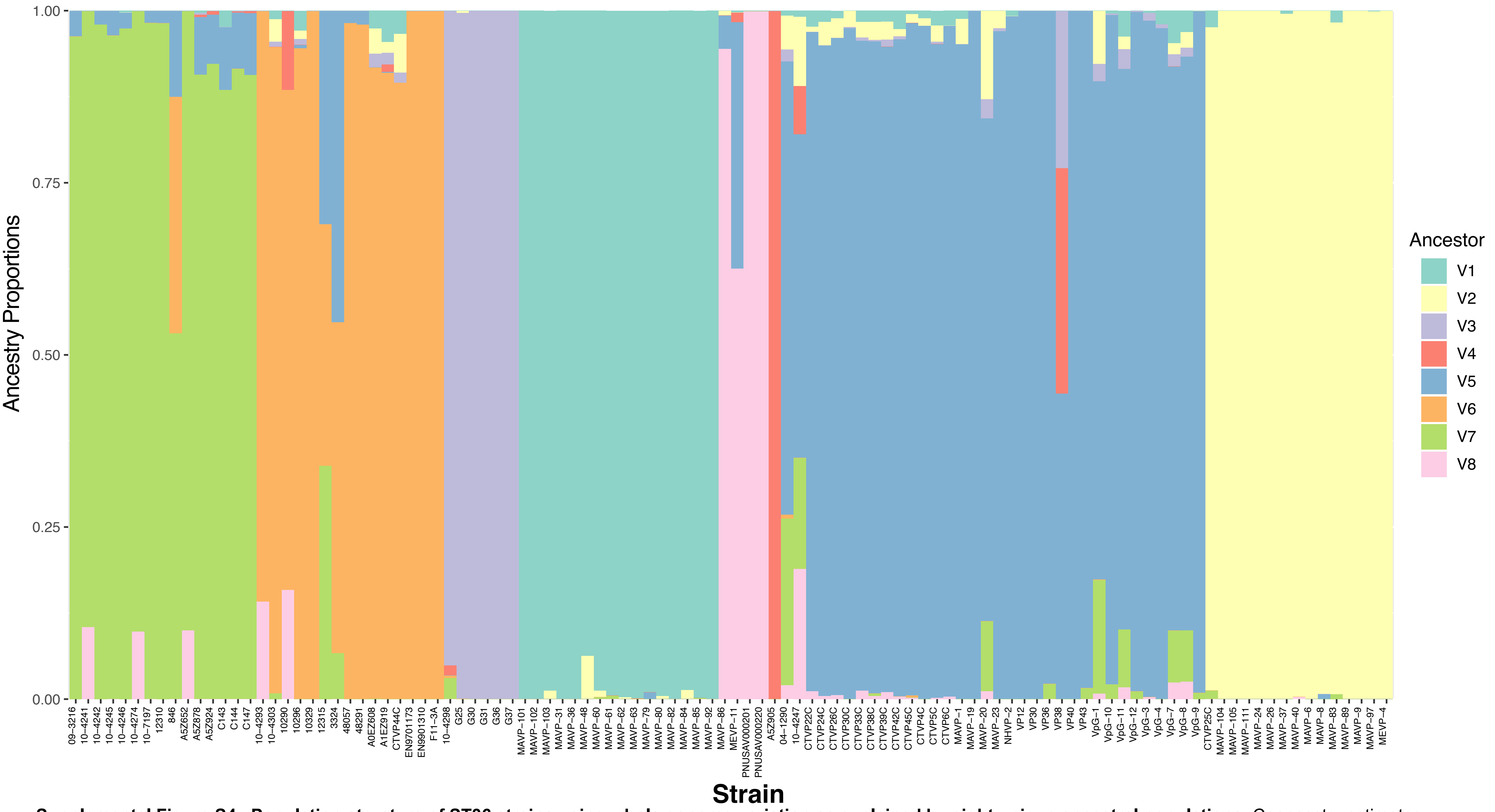


**Supplemental Figure S4. Population structure of ST36 strains using whole genome variation as explained by eight unique ancestral populations.** Coancestry estimates were inferred from core genomes of ST36 strains by LEA (Frichot and François, 2015). Colored bars represent proportions of genetic variation derived from eight ancestral populations which generated the lowest level of cross-entropy (see Fig. S3B).

### Figure S5

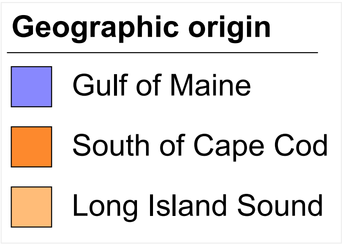


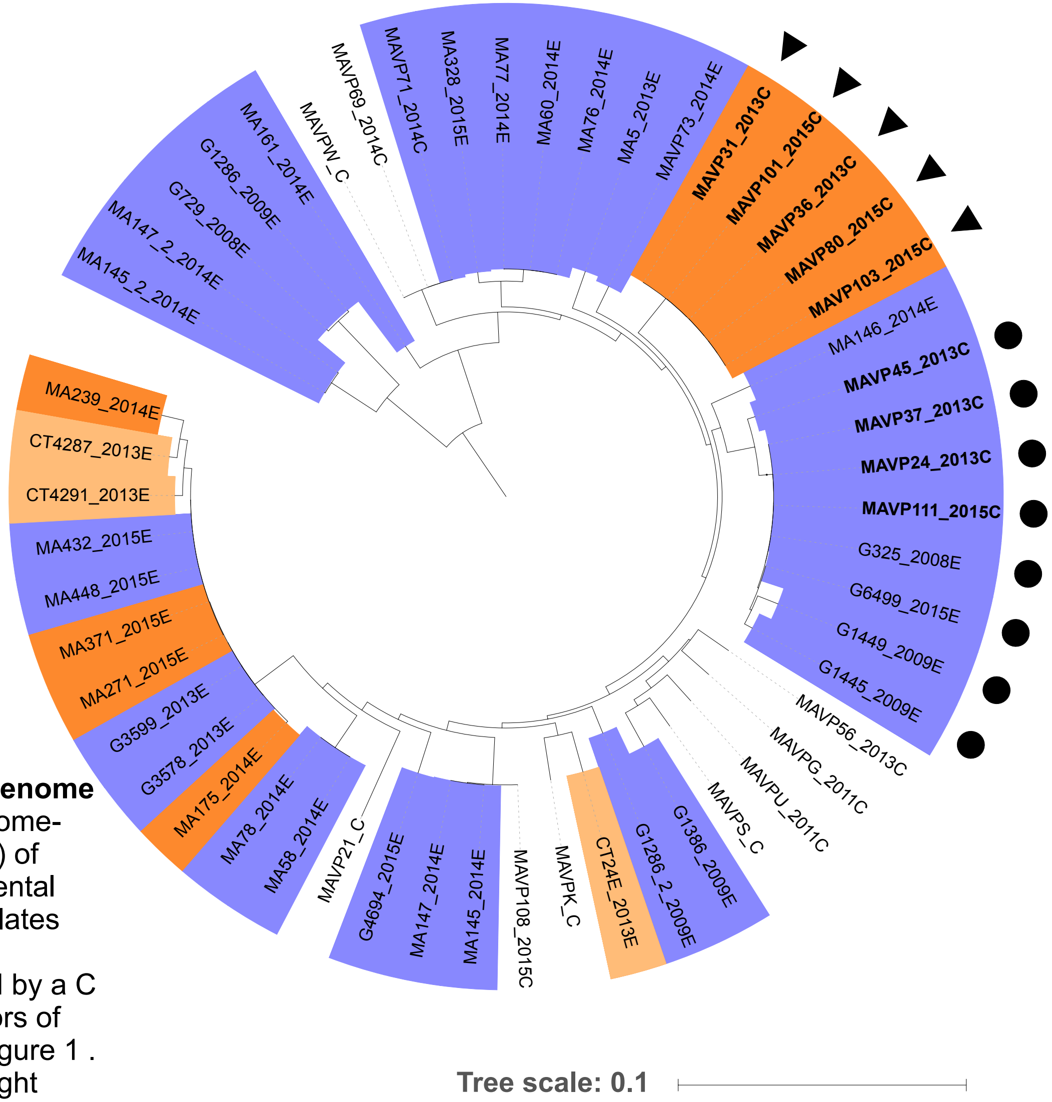


**Supplemental Figure S5. Whole genome ML phylogeny of inoviruses.** Genome-BLAST Distance Phylogeny (GBDP) of whole phage genomes of environmental and clinical *V. parahaemolyticus* isolates with date of isolation. Clinical and environmental isolates are indicated by a C or E after the year of isolation. Colors of geographic traceback match with Figure 1 . Shapes next to isolate names highlight Vipa26 (circle) and Vipa36 (triangle).

**Figure S6**

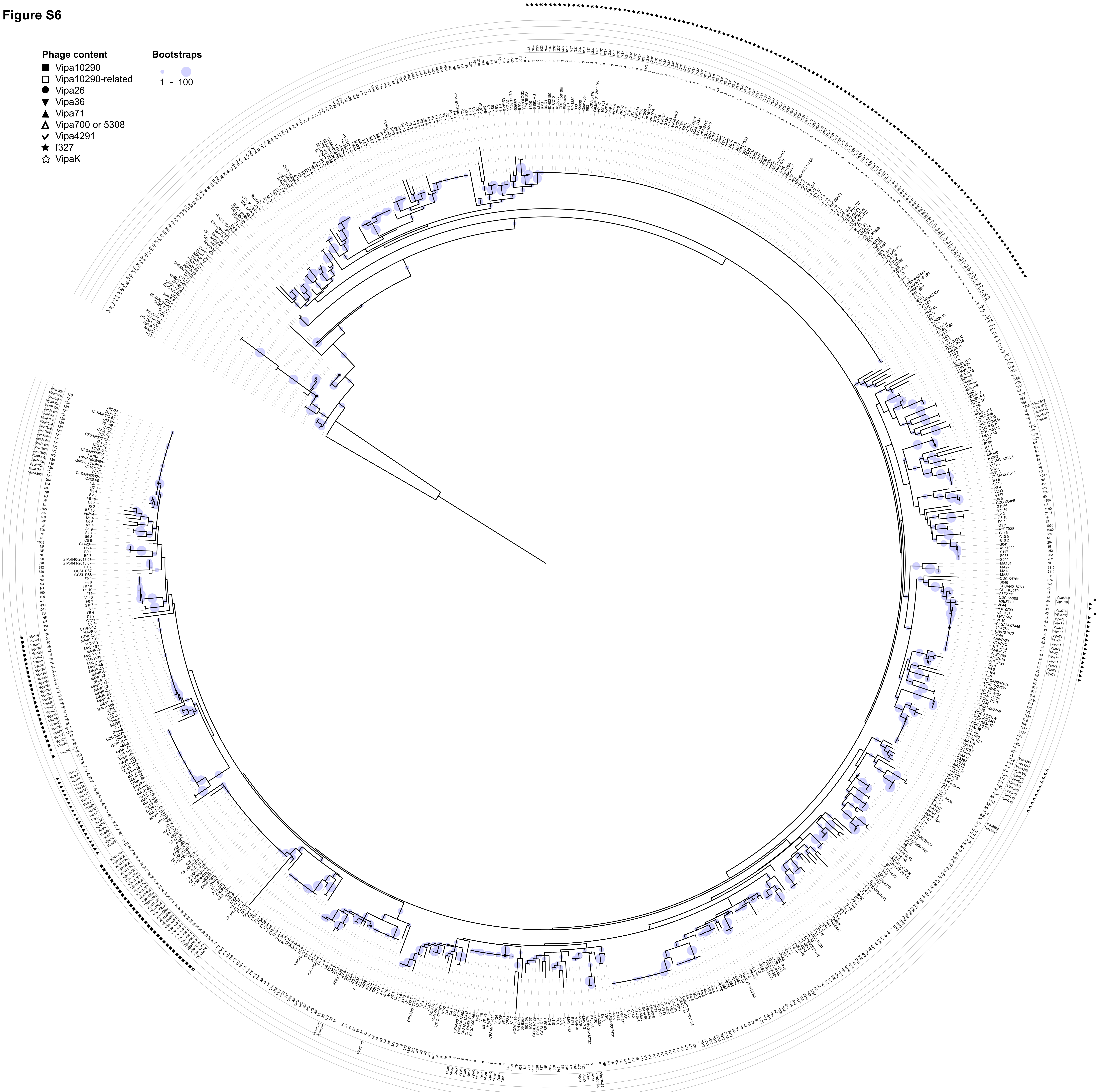


**Supplemental Figure S6**. Maximum-likelihood tree of all core phage in quality *Vibrio parahaemolyticus* genomes. Phage DNA was extracted by alignment using Vipa26 core genome from 736 out of 1500 genomes (Table S3). 1000 bootstraps were mapped onto the best scoring ML tree. Branch ends indicate strain names followed by ST, geographic region, and phage identity (symbol).
